# Human immunodeficiency virus integration complexes are active following ordered addition of wild type integrase, viral DNA, and LEDGF/p75

**DOI:** 10.1101/2022.09.19.508505

**Authors:** Anthony J. Rabe, Jacob A. Russo, Ross C. Larue, Kristine E. Yoder

**Affiliations:** Department of Cancer Biology and Genetics, The Ohio State University College of Medicine, 460 West 12^th^ Ave. Columbus, OH, 43210 USA; Comprehensive Cancer Center, The Ohio State University, Columbus, OH, 43210 USA; Center for Retrovirus Research, The Ohio State University, Columbus, OH, 43210 USA

**Keywords:** Retrovirus, HIV-1, integrase, LEDGF/p75, enzymology

## Abstract

Human immunodeficiency virus (HIV-1) requires integration of the viral genome into the host DNA for replication. Efficient HIV-1 integration employs a host co-factor LEDGF/p75 to stabilize the HIV-1 integration complex and tether that complex to host chromatin. Integration may be studied with purified components HIV-1 integrase (IN), LEDGF/p75, and DNA mimicking the ends of the viral DNA genome (vDNA) assembled as an intasome. There is a likely order of addition during infection with HIV-1 IN binding to vDNA before encountering LEDGF/p75. However, the ordered assembly of wild type HIV-1 IN, LEDGF/p75, and oligomer vDNA has not been tested. Variable assemblies occurred on ice before the addition of target DNA. Incubation on ice and addition of LEDGF/p75 were required to assemble complexes capable of efficient concerted integration. Integration efficiency following variable order of addition of intasome components was greatest when LEDGF/p75 was added last to preassembled HIV-1 IN and vDNA.

## 1. Introduction

Retroviruses, including human immunodeficiency virus (HIV-1), must reverse transcribe their genomic RNA to a double stranded DNA (cDNA) and integrate that viral genome into host DNA [1]. Integration is mediated by the viral enzyme integrase (IN) which has three domains: an amino terminal domain (NTD), a catalytic core domain (CCD), and a carboxyl terminal domain (CTD) [1]. Inhibitors of HIV-1 IN (integrase strand transfer inhibitors, INSTIs) were first approved for clinical use in 2007 [2]. This class of antiretroviral drugs has proven to be highly effective for treating HIV-1 infection [3,4]. However, biochemical and biophysical studies of HIV-1 IN are hampered by the enzyme’s inherent insolubility and propensity to aggregate [5]. Engineering a highly soluble, non-specific DNA binding protein Sso7d from *Sulfolobus solfataricus* to the amino terminus of HIV-1 IN led to increased solubility and integration activity *in vitro* [5,6]. The residues mediating DNA binding were altered to generate Sso7d_mut_ [5]. The chimeric Sso7d_mut_-IN has allowed cryo-EM studies of complexes (intasomes) assembled with DNA oligomers mimicking the viral cDNA ends (vDNA) or DNA oligomers recapitulating vDNA covalently joined to target DNA, as well as complexes with an INSTI [7,8]. It was inferred that the 7 kDa Sso7d solubility domain did not affect the multimerization of HIV-1 IN, but structures without the solubility domain have not been resolved.

Efficient HIV-1 integration during infection requires the host-co-factor lens epithelium derived growth factor (LEDGF/p75) [9,10]. LEDGF/p75 is a transcription activator that includes a chromatin binding PWWP domain at the amino terminus, two AT-hook domains that bind tracts of A and T in DNA, and an HIV-1 IN binding domain (IBD) at the carboxyl terminus [11]. The IBD binds a cleft between CCDs of two HIV-1 IN protomers of the intasome [12]. LEDGF/p75 acts as a tethering factor bringing the HIV-1 IN complex into proximity of select chromatin features [13–16]. Deletion of LEDGF/p75 reduces integration during an infection 10-fold [9,10]. Additionally, this host co-factor has been shown to stabilize HIV-1 intasomes [17]. Sso7d_mut_ chimeric HIV-1 IN forms mostly tetramers with two vDNA oligomers, but in the presence of IBD appears to be mostly dodecamers, suggesting that LEDGF/p75 participates in assembly and/or stability of these complexes [8].

We evaluated integration efficiency of HIV-1 IN without a solubility domain in the presence of full-length LEDGF/p75. The order of addition was varied for the intasome components IN, LEDGF/p75, and vDNA to evaluate optimal assembly and catalytic activity. It seems likely that during an infection HIV-1 IN will interact with nascent viral cDNA ends within a capsid core and later interact with LEDGF/p75 in the nucleus. Yet the intasome is empirically unstable in the absence of LEDGF/p75. Some studies have suggested that LEDGF/p75 is present in the virion where it would interact with HIV-1 IN before generation of vDNA [18]. The order of addition of intasome components with short vDNA and wild type HIV-1 IN has not been systematically tested for the assembly and kinetics of active integration complexes yet has implications for future biochemical and biophysical studies.

## 2. Materials and Methods

### 2.1 Purification of HIV-1 IN and LEDGF/p75

The Sso7d_mut_-IN expression plasmid was modified to include a sortase cleavage site LPETGG upstream of HIV-1 IN separated by a single G residue [5,19]. Recombinant HIV-1 and LEDGF/p75 were purified as previously described [20–22]. Briefly, Sso7d_mut_-sortase cleavage site-IN and LEDGF/p75 were expressed in *E. coli* Rosetta (DE3) pLysS (Millipore Sigma). Expression was induced with IPTG (Gold Biotechnology). Sso7d_mut_-sortase cleavage site-IN was purified with nickel affinity chromatography (Ni-NTA Superflow, Qiagen). The hexahistidine tag and Sso7d_mut_ were cleaved from HIV-1 IN by addition of his-tagged sortase and GGGC peptide (GenScript). The reaction was exposed to nickel resin to remove sortase and any residual uncleaved IN. HIV-1 IN was further fractionated by size exclusion chromatography (SEC) with a Superdex 200 Increase (10/300) column (GE Healthcare) and stored at −80°C (Fig. 1A). LEDGF/p75 with a hexahistidine tag was also purified by nickel affinity chromatography. The hexahistidine tag was removed by cleavage with HRV 3C protease. LEDGF/p75 was fractionated by SEC with a HiLoad 16/60 Superdex-200 and stored at −80°C (Fig. 1A).

**Fig. 1.**
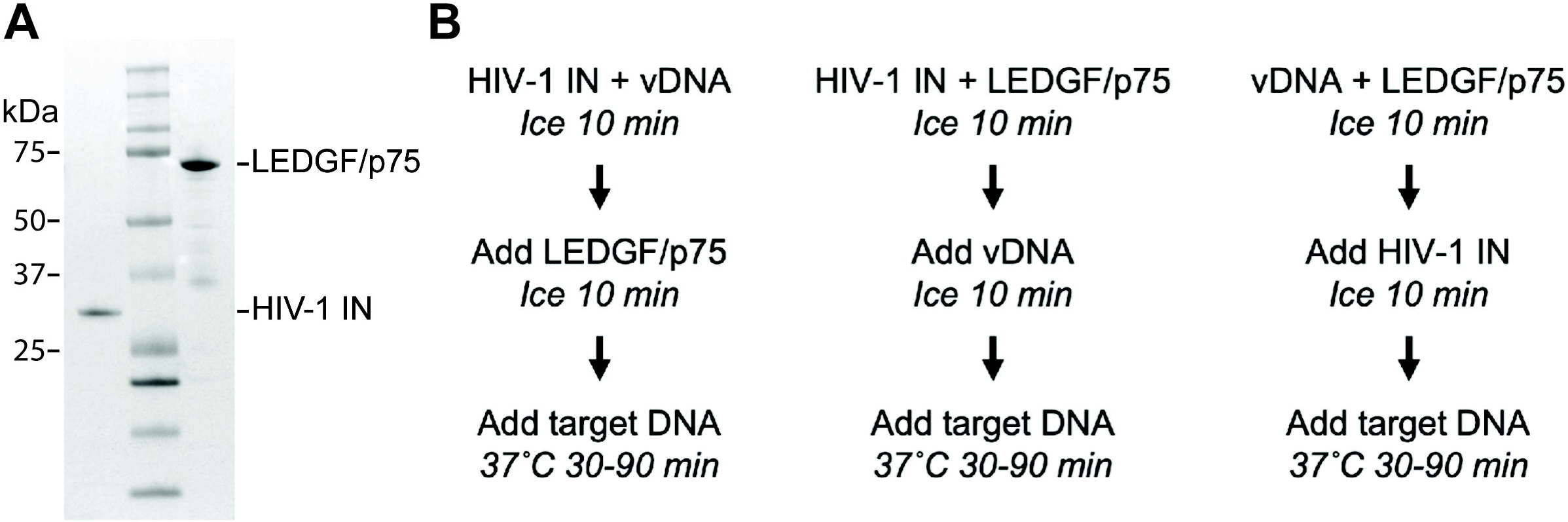
Purified proteins and experimental strategy. **A**) HIV-1 IN and LEDGF/p75 were analyzed by SDS-PAGE stained with Coomassie brilliant blue. Molecular weights are indicated in kDa (left). Lane 1, HIV-1 IN, 32 kDa; Lane 2, protein molecular weight markers; Lane 3, LEDGF/p75, calculated molecular weight 60 kDa, apparent molecular weight approximately 75 kDa. **B**) Three variations of order of addition were tested. The variable reactants - HIV-1 IN, vDNA, and LEDGF/p75 - were diluted in reaction buffer and incubated on ice for 10 min after each addition to allow association. A supercoiled plasmid target DNA was added and samples were immediately transferred to 37°C for 30, 60, or 90 min. Reactions were stopped by the addition of proteinase K, SDS, and EDTA.

### 2.2 Integration reactions

DNA oligomers mimicking the viral cDNA ends (vDNA) U5_25 5’ AGCGTGGGCGGGAAAATCTCTAGCA 3’ and U5_25b 5’ ACTGCTAGAGATTTTCCCGCCCACGCT 3’ (Integrated DNA Technologies, IDT). The oligomer U5_25 was labeled at the 5’ end with a Cy5 fluorophore (IDT). The DNA oligomers were annealed as previously described [23].

Integration assays were performed in 25 mM HEPES, pH 7.5, 110 mM NaCl, 10 mM MgCl_2_, 4 μM ZnCl_2_, 10 mM DTT, 1 mM vDNA, 50 ng pGemT in a total volume of 15 μL. Chemicals were of the highest grade (Sigma Aldrich). Protein concentrations were 2 μM HIV-1 IN and 0.5 μM LEDGF/p75. Reaction components were assembled on ice as indicated in Fig. 1B and transferred to 37°C immediately after addition of supercoiled pGemT target DNA (Promega). Reactions were incubated for 30, 60, or 90 min and stopped by the addition of 0.5% SDS, 0.5 mg/mL proteinase K, 25 mM EDTA, pH 8.0, and incubated at 37°C for 1 h. Reaction products were separated by 1.2% agarose gel electrophoresis in TAE, stained with 0.5 μg/mL ethidium bromide, and imaged for ethidium bromide and Cy5 fluorescence (Sapphire Biomolecular Imager, Azure Biosystems). HSI, CI, and vDNA Cy5 fluorescence values were added to generate a total fluorescence value for each lane; these values were used to calculate HSI and CI products as a percentage of the total fluorescence in each lane, the percent integration (AzureSpot Pro, Azure Biosystems). Error bars indicate the standard deviation between at least three independent experiments. Integration assays were performed with two independent purifications of HIV-1 IN and LEDGF/p75. *p* values were calculated by two tail paired t test (Microsoft Excel).

## 3. Results

### 3.1 Wild type HIV-1 IN and short vDNA require LEDGF/p75 for efficient integration

Recombinant HIV-1 IN and LEDGF/p75 were purified from bacteria (Fig. 1A). DNA oligomers were annealed to generate vDNA and included a single Cy5 fluorophore at the 5’ end distal to IN binding. These two proteins and the vDNA were added to reaction buffer in a variable order of addition on ice and allowed to assemble (Fig. 1B) [24]. Then a 3 kbp supercoiled plasmid was added as the integration target and samples were immediately transferred to 37°C. The integration reactions were incubated for 30, 60, or 90 min and stopped by the addition of proteinase K, SDS, and EDTA. The samples were incubated at 37°C for an additional hour. Reaction products were separated by 1.2% agarose gel electrophoresis and visualized by ethidium bromide and Cy5 fluorescence imaging (Fig. 2B). Concerted integration (CI) of two vDNAs to the target results in a 3 kbp linear DNA while half site integration (HSI) of a single vDNA to the target results in a tagged circle (Fig. 2A). While vDNA tagged circles have the same mobility as a relaxed circle, Cy5 fluorescence distinguishes HSI products from relaxed plasmids resulting from non-specific nuclease activity [25,26]. The Cy5 fluorescence of HSI, CI, and vDNA bands were quantified and used to calculate integration as the percentage of total fluorescence in each lane.

**Fig. 2.**
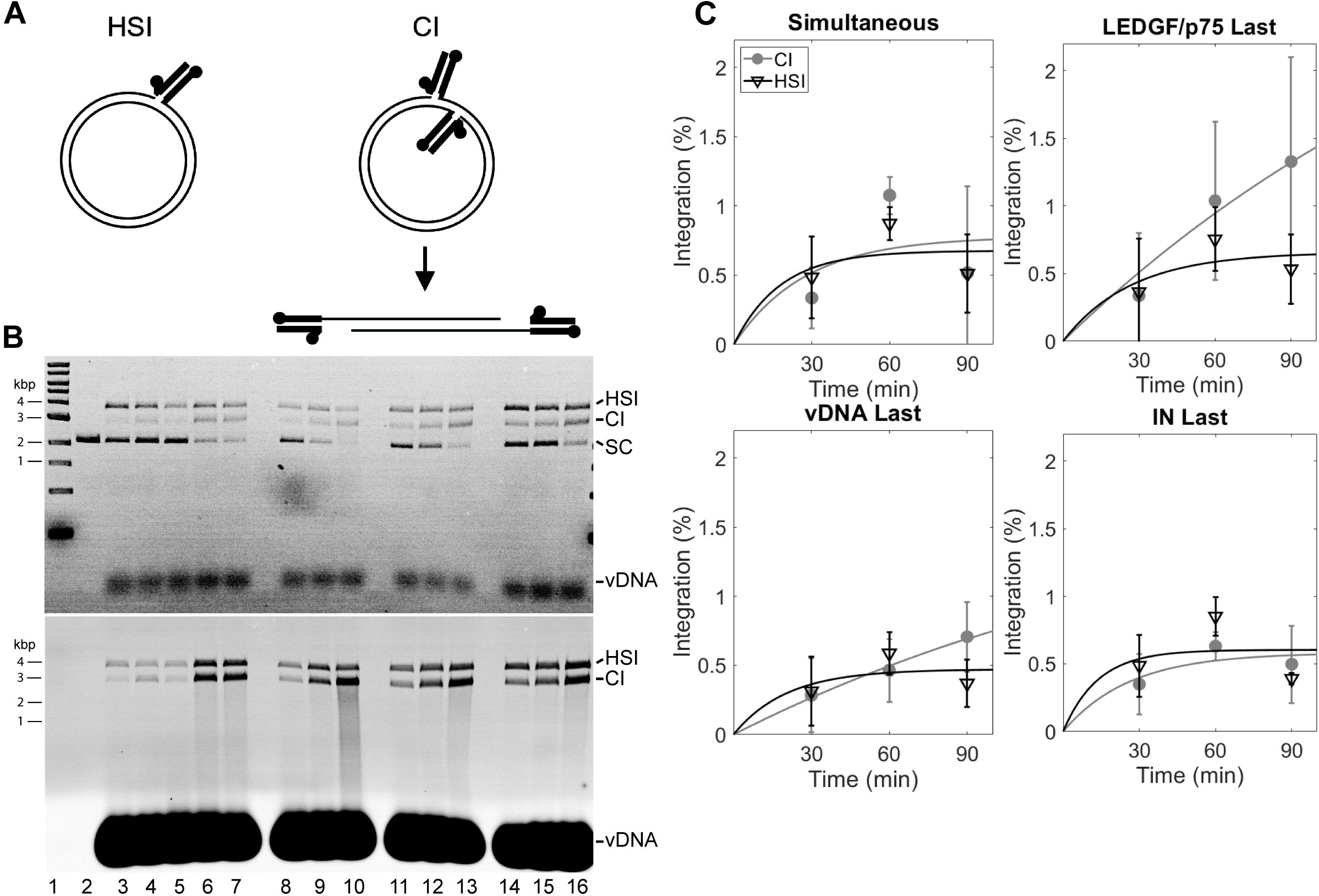
HIV-1 integration is most efficient following ordered assembly on ice with LEDGF/p75 added last. **A**) Two products result from integration *in vitro* with an oligomer vDNA (heavy black lines) and supercoiled plasmid target DNA (light black lines) substrates. Half site integration (HSI) products have a single vDNA covalently joined to the plasmid DNA target. This integration introduces a nick in the plasmid leading to a relaxed circle. Concerted integration (CI) mimics the physiologically relevant integration reaction with two vDNAs joined to the plasmid DNA 5 bp apart. This DNA melts to a linear product flanked by vDNA. Filled circles indicate 5’ ends. **B**) HIV-1 IN, Cy5 labeled vDNA, and LEDGF/p75 were added to integration reactions with variable order of addition or simultaneously. Following assembly of complexes on ice, 50 ng supercoiled plasmid (SC) was added and reactions were incubated at 37°C for 30, 60, or 90 min. CI products have the mobility of a 3 kbp linear DNA, HSI products have the mobility of a relaxed plasmid and migrate slower than CI products. Reaction products were resolved by agarose gel electrophoresis and visualized by ethidium bromide staining (top) and Cy5 fluorescence (bottom). The Cy5 image allows discernment of fluorescent HSI products and unlabeled nicked plasmid. DNA sizes are indicated in kbp (left). Lane 1, DNA ladder; Lane 2, supercoiled plasmid target DNA; Lane 3, HIV-1 IN and vDNA incubated on ice 20 min before addition of target DNA; Lane 4, HIV-1 IN, LEDGF/p75, and vDNA added to target DNA without incubation on ice; Lanes 5-7, HIV-1 IN, LEDGF/p75, and vDNA combined simultaneously and incubated on ice 20 min before addition of target DNA and incubation at 37°C; Lanes 8-10, order of addition with LEDGF/p75 added last; Lanes 11-13, order of addition with vDNA added last; Lanes 14-16, order of addition with HIV-1 IN added last. Three lanes correspond to incubation at 37°C with target DNA for 30, 60, and 90 min. **C**) HSI, CI, and vDNA were quantified by Cy5 fluorescence. HSI (open triangles) and CI (filled circles) products are expressed as the percentage of the total fluorescence in the lane (Integration (%)). Error bars indicate the standard deviation between three independent experiments.

HIV-1 IN and vDNA were incubated on ice for 20 min followed by addition of supercoiled plasmid DNA and incubation at 37°C for 90 min. In these reactions that did not include LEDGF/p75, we observed HSI products (Fig. 2B, Lane 3, HSI = 0.20 ± 0.15% integration). It has previously been shown that HSI products may be catalyzed by HIV-1 IN dimers that do not include LEDGF/p75 [27]. CI products were discernable but could not be quantified above background (Fig. 2B, Lane 3, CI = 0.15 ± 0.16% integration). The dearth of linear products without LEDGF/p75 confirms the necessity of this protein for efficient CI either due to effects on assembly or stability of the intasomes.

### 3.2 Assembly of intasome components is more efficient on ice than at 37°C

The simultaneous addition of all intasome components with and without incubation on ice was compared. Limited integration was observed without incubation on ice (Fig. 2B, Lane 4, HSI = 0.28 ± 0.03%, CI = 0.22 ± .014%). When reaction components were incubated on ice for 20 min before the addition of target DNA, CI and HSI products were readily apparent at all time points (Fig. 2B, Lanes 5-7). The total HSI and CI products were not significantly different at 90 min (p > 0.05), but were greater (Fig. 2B, Lane 7, HSI = 0.51 ± 0.28%, CI = 0.52 ± 0.63%) than observed without incubation on ice. This observation indicated the necessity for assembly on ice before addition of target DNA and transfer to 37°C for integration.

### 3.3 The most efficient order of addition is LEDGF/p75 added last

Three variations of order of addition were assayed (Fig. 1B). Reactions with two components were incubated on ice for 10 min followed by addition of the third component and further incubation on ice for 10 min before initiation of the integration assay. The addition of LEDGF/p75 last yielded the greatest CI products at 90 min (Fig. 2B, Lane 10, CI = 1.33 ± .053%). A smear of additional CI events into the HSI and CI products is present and has faster mobility than CI products, indicating highly active intasomes. HSI and CI products were also apparent with vDNA added last (Fig. 2B, Lanes 11-13) or HIV-1 IN added last (Fig. 2B, Lanes 14-16), but these orders of addition displayed the least CI activity. With all orders of addition HSI products appeared early without much accumulation over time. Similarly, CI products did not appear to accumulate dramatically over time, except when LEDGF/p75 was added last. Together these data indicate that HIV-1 integration complexes were most active following an ordered assembly on ice with LEDGF/p75 added last.

## 4. Discussion

HIV-1 IN is present in the virion before reverse transcription occurs. The order of assembly of intasome components during infection may be interaction of HIV-1 IN with nascent vDNA followed by LEDGF/p75 binding in the nucleus. Late association of LEDGF/p75 would then stabilize the intasome and mediate binding to target DNAs via the PWWP or AT hook domains. Alternatively, it is possible that a partially assembled HIV-1 IN and vDNA complex requires LEDGF/p75 for formation of the complete intasome complex in the nucleus. Some reports have suggested LEDGF/p75 is packaged within the virion leading to HIV-1 IN interaction with its host co-factor before reverse transcription [18]. Intuitively the efficient assembly of HIV-1 intasomes *in vitro* would follow a similar order of addition as during infection. Deviation from the natural order of addition during assembly *in vitro* could result in conflicting results. These results are informative for future structural and functional studies of HIV-1 intasomes.

In the absence of LEDGF/p75 we detected HSI products and very little CI. Previous similar studies of HIV-1 IN integration showed significant CI and HSI in the absence of LEDGF/p75 [28,29]. These studies employed longer viral donor DNAs ranging in length from 0.48 to 4.1 kbp. Long viral donor DNAs display greater activity in integration assays *in vitro* compared to oligomer vDNA. Donor vDNAs of several hundred bp have been suggested to form a stable synaptic complex by allowing the binding of multiple IN protomers which cannot be accommodated by a 25 bp vDNA [30]. In contrast, HIV-1 IN forms dimers in solution which can bind short vDNA oligomers and perform HSI [31,32]. Integration by Sso7d_mut_-IN with short vDNA oligomers displays more HSI than CI products [8]. Shorter vDNA donors, like those used in this study, that are known to lead to less integration, are more commonly used in structural studies of intasomes [7,8]. Ideally wild type intasomes with short vDNA capable of efficient CI would be generated for such studies.

A previous study of order of addition with long vDNA observed greater CI than HSI when LEDGF/p75 was added to HIV-1 IN bound to vDNA [28]. This current study with short vDNA recapitulates the observation of greater CI when LEDGF/p75 is added last. Interestingly, the ordered addition of intasome components led to greater integration than simultaneous addition of wild type HIV-1 IN, short vDNA, and LEDGF/p75. Further, we found that incubation on ice to allow assembly of intasome components before addition of target DNA was necessary for efficient integration. The lack of incubation on ice was nearly as detrimental to CI as lack of LEDGF/p75. This suggests that the intasome components were better able to assemble on ice and did not assemble well when incubated at 37°C. It is possible that the ice incubation recapitulates the slow addition of intasome components similar to infection when a capsid core containing HIV-1 IN and vDNA must traverse the cytoplasm and enter the nucleus with a possible gradual addition of LEDGF/p75 to the nascent intasome.

Taken together, these data suggest that wild type HIV-1 IN without a solubility domain may be assembled with short vDNA and LEDGF/p75 into functional intasomes. The assembly conditions here may be adapted to larger volumes for purification of unmodified intasomes to more closely resemble infection conditions. These complexes may better recapitulate physiological integration complexes for biochemical and biophysical experiments. Furthermore, these studies highlight the importance of ordered assembly for functional intasomes during HIV-1 infection suggesting the potential for new antiretroviral therapeutic targets. Indeed, the search for novel HIV-1 IN targeting factors with unexploited modes of action is an ongoing area of interest to combat resistance to current therapies [33,34].

## Acknowledgements

This work was supported by NIH AI150496 to KEY.

